# Machine learning classification of fibromyalgia from blood transcriptomes: confounder benchmarking and implications for endometriosis-associated pain

**DOI:** 10.64898/2026.10.21.753755

**Authors:** Salsabil Ouni, Ferdaous Ouni, Alif Chebbi

## Abstract

Fibromyalgia (FM) is a chronic nociplastic pain disorder characterized by widespread musculoskeletal pain, profound fatigue, sleep disturbances, and cognitive dysfunction. Driven by central sensitization, it involves aberrant neuroplastic amplification of central nociceptive processing without structural or inflammatory tissue damage. It lacks an objective biomarker, and blood transcriptomic signatures have been proposed to fill this gap. This study reanalysed the largest public peripheral blood mononuclear cell (PBMC) RNA-seq cohort of FM (96 patients, 93 controls) using leakage-free nested cross-validation. Gene expression discriminated patients from controls (AUC 0.82), well above a permutation null. However, two disease-agnostic baselines approached this performance: sex alone (AUC 0.73), reflecting a 95% versus 44% female imbalance between groups, and a four-cell-type composition proxy (AUC 0.71–0.76). In females only, expression retained discriminative signal (AUC 0.78) but did not significantly exceed composition alone. Interferon and lysosomal gene sets were depleted in FM, but this depletion was lost after adjusting for lower monocyte content. These results indicate that much of the apparent PBMC signature of FM reflects sex and cell composition, and that blood-based classifiers for chronic pain, including related nociplastic conditions such as endometriosis, should be benchmarked against these confounders before any clinical use.

Figure 1:
Graphical Abstract.

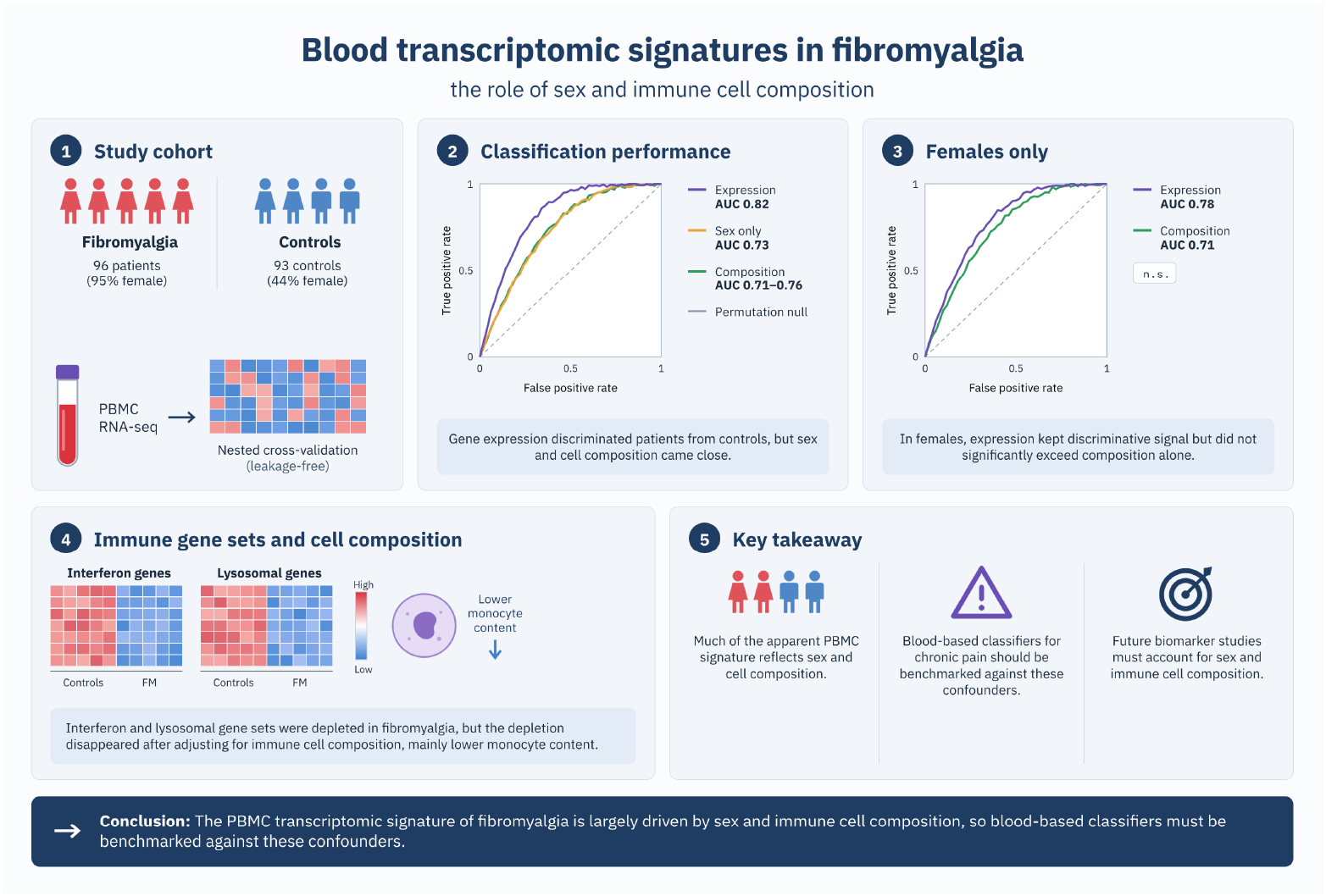

## 1 Introduction

Fibromyalgia (FM) is a prototypical nociplastic pain condition: widespread musculoskeletal pain, persistent fatigue, unrefreshing sleep, and cognitive difficulties arise from a disturbance of central pain processing itself, rather than from any identifiable tissue injury or inflammatory lesion [1]. It is diagnosed clinically, on the basis of these symptoms meeting the 2016 ACR criteria [2], in the absence of any validated laboratory or imaging biomarker. Its proposed biological substrates include central sensitization (amplified central nervous system signaling producing hyperalgesia and allodynia), impaired descending pain-inhibitory control, and neuroimmune and glial activation, rather than a single peripheral lesion or structural abnormality. Population prevalence estimates vary substantially by methodology and diagnostic criteria (individual studies range 0.2–6.6%); a large meta-analysis pooling *>* 3.6 million subjects across 65 studies estimated general-population prevalence at 1.78% (95% CI 1.65–1.92) [3].

The nociplastic pain framework extends beyond FM to other chronic pain conditions that share its central-sensitization mechanisms, among them endometriosis, in which nociceptive, neuropathic, and nociplastic contributions to pain can coexist [4]. FM and endometriosis co-occur more often than expected by chance: in a comparative cohort of 668 women with endometriosis, 7% also met FM diagnostic criteria, and this subgroup showed more severe pelvic pain and a higher burden of autoimmune and psychiatric comorbidity than endometriosis alone [5]. Both conditions are hypothesized to share peripheral neuroimmune features, including altered circulating immune cell composition and interferon-pathway activity. Consequently, any blood-based classifier for FM requires the same scrutiny of immune cell and demographic confounders that would be expected in endometriosis research.

This diagnostic gap and shared-mechanism rationale have motivated a series of transcriptomic and proteomic profiling studies aimed at identifying molecular correlates of FM. Mohapatra et al. (2024) performed RNA sequencing on PBMCs from 96 FM patients and 93 matched controls. Hierarchical clustering reassigned 5 samples relative to their clinical labels (2 FM patients toward the control profile, 3 controls toward the FM profile); these 5 were dropped, leaving 94 FM and 90 control samples for downstream analysis. Within that 94-patient FM cohort, principal component analysis and hierarchical clustering identified three expression subgroups, with 4 additional patients left unclassified as outliers: FM1 (*n* = 43, extracellular matrix organization and downregulated RhoGDI signaling), FM2 (*n* = 30, reduced inflammatory mediator expression with increased CLEAR/lysosomal-biogenesis gene expression), and a combined FM3/FM4 group (*n* = 17, split into subclusters of 8 and 9 patients respectively, interferon alpha/beta and JAK-STAT signaling together with death-receptor and necroptosis pathway activity) [6]. That study’s contribution is descriptive rather than predictive: it characterizes subtype heterogeneity within FM patients using unsupervised methods, but it does not report a cross-validated classifier, a permutation baseline, or any test of whether its subgroup-defining genes would also emerge from a supervised, whole-cohort comparison against controls.

With roughly 189 samples and 22,000 candidate genes, the main methodological hazard for any supervised reanalysis of this dataset is leakage: running feature selection or other data-dependent preprocessing on the full dataset before cross-validation, which inflates apparent performance in a way that is easily overlooked [7, 8]. A second hazard specific to this cohort is confounding by patient sex: FM is disproportionately diagnosed in women in clinically-referred samples, a pattern partly attributable to referral and diagnostic bias [9], and, per the endometriosis comorbidity rationale above, potentially compounded by gynecological and hormonal factors not captured by sex alone. These are treated here as questions to be tested directly rather than caveats to mention in passing, with results reported regardless of outcome. The present study therefore reanalysed the same public dataset (GEO accession GSE221921) with three objectives: (1) to estimate, without leakage, the ability of gene expression to discriminate FM from controls, and to benchmark this performance against simple baselines based on sex and PBMC composition; (2) to identify the genes driving this discrimination and assess their stability; and (3) to determine whether these genes correspond to the pathways of the original FM subtypes or are better explained by differences in cell composition. The machine learning and statistical terms used throughout are defined in Box 1.

### Box 1. Glossary of machine learning and statistical terms

| Term | Definition as used in this study |
| --- | --- |
| Classifier | A statistical model that assigns each sample to a class (here, FM or control) on the basis of its gene expression profile. |
| Training and test sets | Subsets of samples used, respectively, to fit the model and to evaluate it. Test samples are never used during model fitting. |
| Cross-validation | A resampling procedure in which the data are partitioned into $k$ subsets (here, $k = 5$); each subset serves once as the test set while the remaining subsets are used for training, so that every sample is evaluated exactly once. |
| Nested cross-validation | A second, inner cross-validation performed within each training set to select model parameters (for example, the number of genes) without access to the test samples. |
| Data leakage | Any use of information from test samples during model construction, such as gene selection on the complete dataset. Leakage leads to overestimated performance. |
| Overfitting | The fitting of noise specific to the training samples, resulting in poor performance on new samples. |
| ROC-AUC (AUC) | The area under the receiver operating characteristic curve; equivalent to the probability that a randomly selected FM patient receives a higher predicted score than a randomly selected control. A value of 0.5 corresponds to chance and 1.0 to perfect discrimination. |
| Permutation test | Repetition of the entire analysis with randomly shuffled diagnosis labels, providing the distribution of performance expected by chance alone. |
| Baseline model | A deliberately simple reference model (for example, sex only) that an informative gene-expression model should outperform. |
| Logistic regression; XGBoost | Two classifier families: logistic regression models the outcome as a weighted linear combination of genes, whereas XGBoost (gradient boosting) combines a large number of shallow decision trees. |
| Regularization | A penalty on model complexity that limits overfitting. |
| SHAP values | SHapley Additive exPlanations: a measure of the contribution of each gene to the prediction for each sample. |
| Calibration | The agreement between predicted probabilities and observed frequencies; for instance, samples assigned a 70% probability of FM should be FM in approximately 70% of cases. |
| GSEA | Gene set enrichment analysis: a test of whether the genes of a predefined pathway are concentrated at the top or bottom of a ranked gene list. |

## 2 Methods

The analytical workflow is summarized in Figure 2. It comprises data acquisition and preprocessing, classifier training and evaluation by nested cross-validation, validity and confounder checks, and biological interpretation by explainability and gene set enrichment analyses.

**Figure 2:**
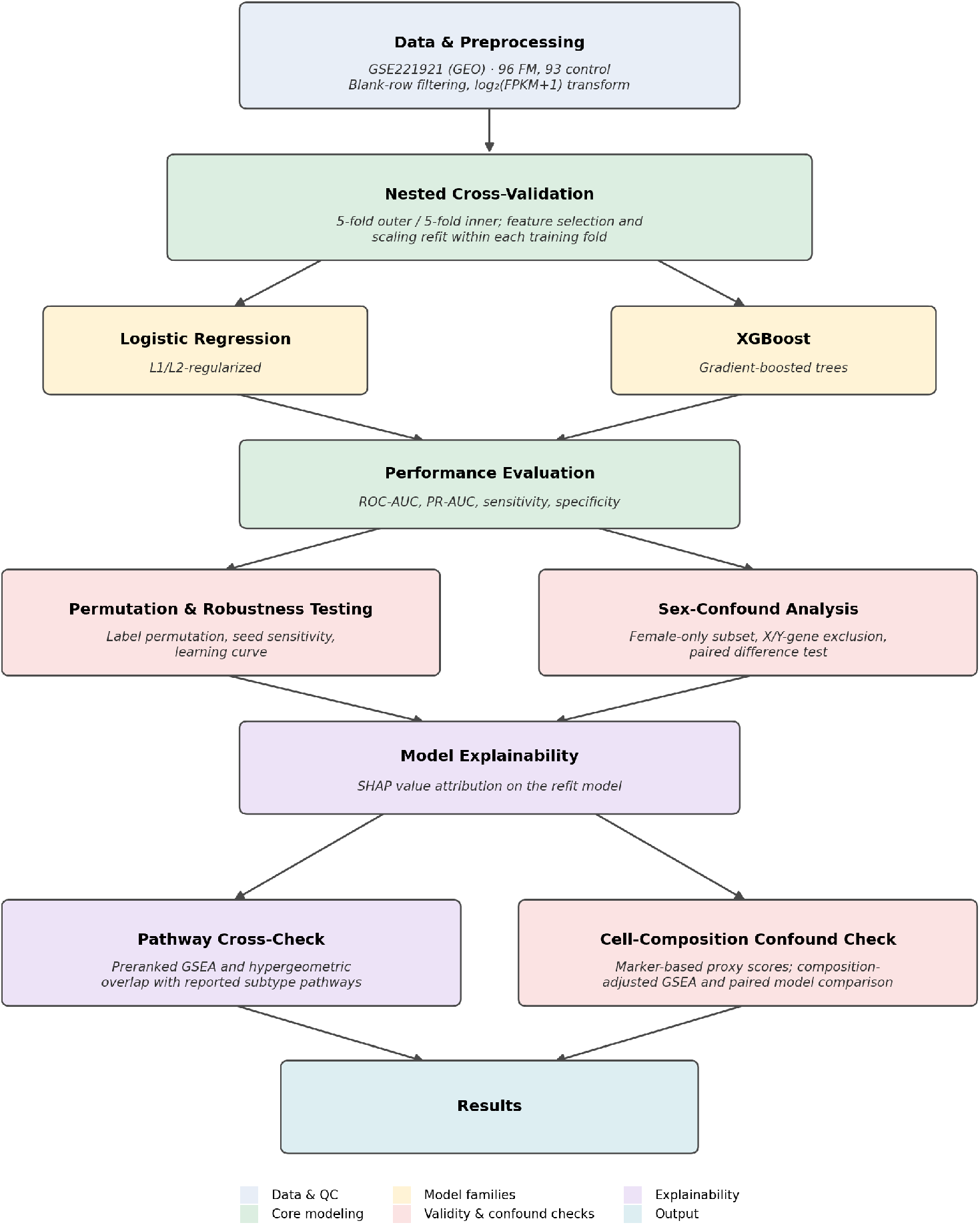
Schematic overview of the analysis pipeline, from the public data to the validity, confounder and robustness checks and the biological interpretation.

### 2.1 Data

The author-processed FPKM (fragments per kilobase of transcript per million mapped reads) expression matrix and sample metadata for GSE221921 were downloaded from the NCBI GEO supplementary spreadsheet and series matrix file, with dates and checksums recorded in the code repository [12]; no raw FASTQ reprocessing was needed. The dataset comprises 189 samples (96 FM, 93 control), matching the source publication. Sex is available for all samples and agreed across the two GEO metadata sources; age and sequencing batch are not in the public metadata (the source paper reports a median age of 48 vs. 45 [control] in its text). All sample-level metadata used in this reanalysis, sample ID, FM/control label, sex, and GEO sample accession for all 189 samples, is released as a single file in the code repository (Data and Code Availability). Gene-level metadata (Ensembl ID, Hugo symbol, chromosome, genomic coordinates) and the biotype annotation used for the protein-coding-only check (Section 2.5) are released alongside it, so none of the metadata relevant to this analysis is confined to prose description. Separately, a parsing artifact in the source spreadsheet was identified and excluded: of 103,810 rows, 81,895 were blank, an inflated sheet dimension rather than missing measurements, leaving 21,915 genes with FPKM values for all samples.

### 2.2 Preprocessing and quality control

No genes required removal for near-zero expression (FPKM = 0 in *>* 90% of samples), consistent with the released matrix already being restricted to detectably-expressed genes. Expression values were log_2_(FPKM + 1) transformed. A PCA of the full, log-transformed matrix was used as a pre-modeling sanity check for gross technical artifacts (e.g. batch-like clustering); it showed partial, non-separable overlap between FM and control along the first two components and no dominant clustering by sex (Figure 3). Sex is a biological and demographic covariate of interest in this cohort, addressed directly in Section 3.3, not a technical confound; its absence from the first two principal components here only indicates that it is not the dominant axis of global transcriptome-wide variance, and does not rule out the smaller, targeted sex-correlated signal characterized later. The incomplete separation between groups is consistent with the transcriptional heterogeneity of FM reported in the source study.

**Figure 3:**
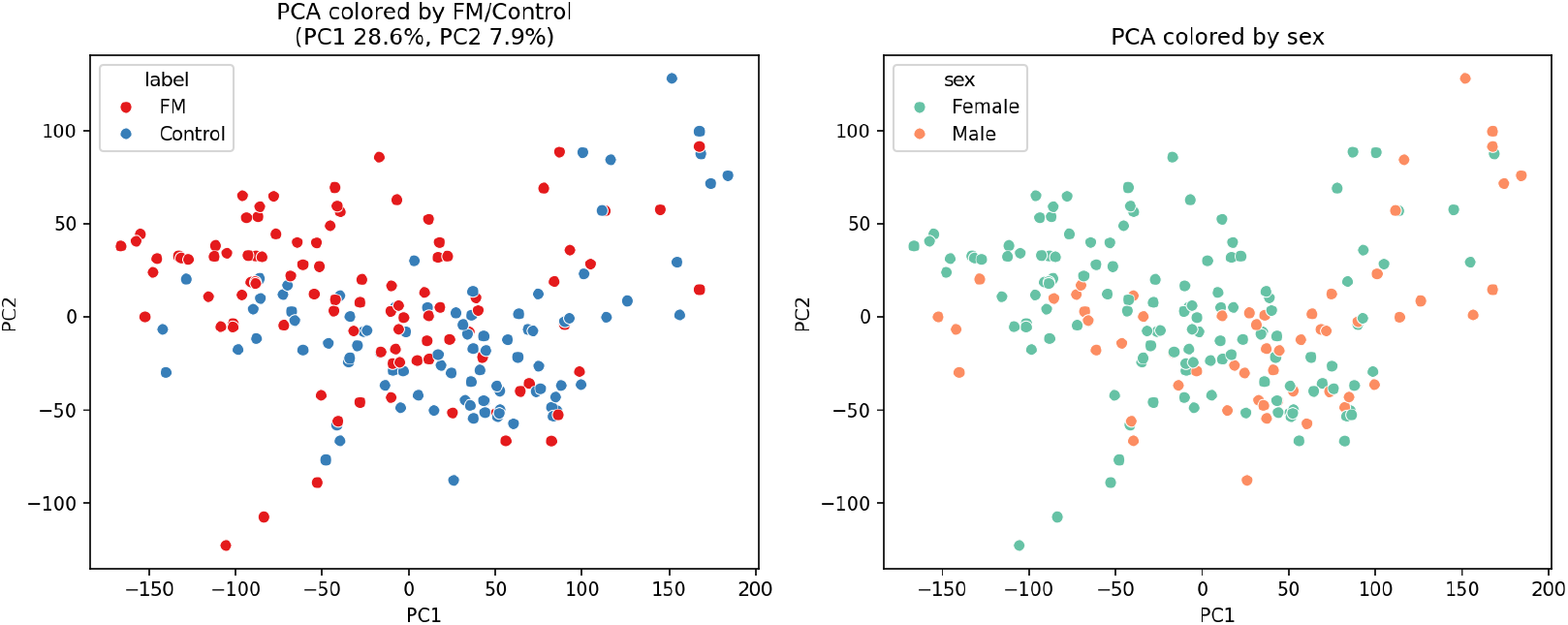
Principal component analysis of log_2_(FPKM + 1) expression. Each point represents one sample, colored by diagnosis (left) or sex (right). Neither variable separates the samples along the first two principal components.

### 2.3 Nested cross-validation

All feature selection and standardization were performed strictly inside a scikit-learn Pipeline (SelectKBest → StandardScaler → classifier), refit from scratch on each training fold by construction, so that no information from a held-out fold could influence feature selection or scaling for that fold. Gene ranking used a vectorized Mann-Whitney U score. Outer evaluation used 5-fold stratified cross-validation; hyperparameters, including the number of selected features *k* ∈ {50, 100, 200}, were tuned via an inner 5-fold stratified grid search (GridSearchCV, scoring = ROC-AUC) on the training portion of each outer fold only. Two model families were compared: L1/L2-regularized logistic regression (*C* ∈ {0.01, 0.1, 1, 10}) and XGBoost gradient-boosted trees (max_depth ∈ {2, 3}, learning_rate ∈ {0.05, 0.1}, n_estimators ∈ {100, 200}) [11]. All random states (numpy, sklearn splitters, XGBoost) were fixed to seed 42.

Performance was summarized across outer folds as ROC-AUC, PR-AUC, and sensitivity, specificity, and accuracy at a 0.5 probability threshold, with 95% confidence intervals from 2,000 bootstrap resamples of the pooled out-of-fold predictions. The two model families were compared with a paired bootstrap on their shared out-of-fold predictions.

### 2.4 Permutation-label null baseline

The entire nested-CV pipeline (feature selection inside each fold, inner-loop hyperparameter search, outer-fold evaluation) was rerun 100 times with the FM/control labels randomly permuted, using a reduced hyperparameter grid for tractability (inner 3-fold CV; select k ∈ {50, 200}, *C* ∈ {0.1, 1}, L2 penalty only). This establishes the null distribution of achievable AUC under the same *n* ≪ *p* feature-selection procedure with no true label signal, which is the single most informative sanity check for a classifier of this shape [7, 8].

### 2.5 Explainability

The better-performing model family by nested-CV AUC was refit on the *full* dataset, with its own hyperparameters re-tuned by a single round of 5-fold CV on the full data, strictly for interpretation. This refit’s own internal CV score is not used as a performance estimate, since by this point the model has seen all outer test folds from Section 2.3. SHAP (SHapley Additive exPlanations) values were computed for the selected features [10] (TreeExplainer for XGBoost, LinearExplainer for logistic regression), and genes were ranked by mean absolute SHAP value.

### 2.6 Biological cross-check

This check tests whether three gene sets corresponding to the original paper’s FM subtypes are coordinately shifted between FM and control across the full transcriptome: FM1 (extracellular matrix/RhoGDI/collagen/GP6 signaling), FM2 (CLEAR/lysosomal biogenesis), and FM3 (interferon/JAK-STAT/death-receptor signaling, the original FM3/FM4 group). The sets are curated proxies for these signatures, not the original study’s own gene lists. FM1 (35 genes) covers extracellular-matrix structural and collagen genes plus RhoGDI/Rho-GTPase and platelet-adhesion components; FM2 (23 genes) covers lysosomal and CLEAR-network genes (cathepsins, vacuolar ATPase subunits, *TFEB*, *MCOLN1* ); FM3 (30 genes) covers type-I interferon response, JAK-STAT signaling, and death-receptor/necroptosis genes, the last of these drawn from the specific example genes the original paper names for its FM3/FM4 group. All three sets were drawn from the literature and fixed before the enrichment analysis was run, and were not revised afterward: one top-30 SHAP gene, *COL6A6*, is thematically related to the FM1 set (which already includes *COL6A1* /*COL6A2* /*COL6A3* ) but was deliberately not added after seeing the classifier’s output, to avoid hindsight bias. The complete gene lists are given in Supplementary Table S5. The primary test was preranked gene set enrichment analysis (GSEA) [13, 14]. Of 21,915 genes, 21,446 have a Hugo symbol, and 72 rows sharing a symbol with another gene (47 duplicated symbols) were collapsed to the most extreme, leaving *n* = 21,374 genes ranked by sign(*̄x*_FM_ − *̄x*_Control_) × − log_10_(*p*) from a Mann-Whitney U test of FM versus control on the full dataset (descriptive, not a performance estimate). Enrichment was tested against 1,000 gene-set permutations (minimum set size 1, as the sets are small). The analysis was repeated in females only and after removing four marker-based composition scores from expression (Supplementary Table S6). A secondary hypergeometric test of overlap between the top-*N* SHAP genes (*N* = 20, 30, 50, 100) and the same gene sets is reported in the Supplementary Materials.

### 2.7 Robustness and validity checks

Eight additional checks were run (Table 1). Six rerun a nested-CV design under one main changed condition (the stability and learning-curve checks use logistic regression only, and the learning curve a smaller grid); calibration reanalyzes stored predictions; the composition check uses simple models.

**Table 1:** Robustness and confounder analyses.

| Analysis | Question addressed | Approach |
| --- | --- | --- |
| Feature-selection stability | Are the same genes selected across data partitions? | 5-fold cross-validation repeated 10 times (50 partitions), logistic regression |
| Univariate test | Does the result depend on the Mann-Whitney test? | Replacement by Welch’s $t$-test and by a moderated $t$-test implementing the empirical Bayes method of limma [15] (in-house implementation, validated on synthetic data) |
| Sex: cohort level | Is the signal attributable to sex? | Sex-only baseline model; complete pipeline restricted to females (91 FM, 41 controls) |
| Sex: gene level | Do sex-chromosome genes drive the signal? | Complete pipeline after exclusion of all 851 X- and Y-linked genes |
| Calibration | Are predicted probabilities reliable? | Brier scores and reliability curves from out-of-fold predictions |
| Learning curve | Would larger samples improve performance? | Training sets subsampled to 30, 50, 70, 85, and 100% (test sets fixed), 20 repetitions each |
| Cell composition | Does the signal reflect differences in PBMC composition? | Monocyte, T-, B-, and NK-cell scores (mean standardized expression of 20 marker genes), compared between groups, used alone as a classifier, and regressed out before the female-only GSEA; a coarse proxy rather than a formal deconvolution |
| Protein-coding genes | Do pseudogenes and noncoding transcripts drive the signal? | Candidate pool restricted to the 16,834 protein-coding genes annotated in Ensembl |

No multiple-comparison correction was applied across these checks, as each addresses a distinct validity question; the permutation nulls remain significant under Bonferroni correction (adjusted *p* = 0.030) and the enrichment tests are FDR-adjusted.

### 2.8 Software and reproducibility

The software environment and the reproducibility verification are summarized in Table 2.

**Table 2:**
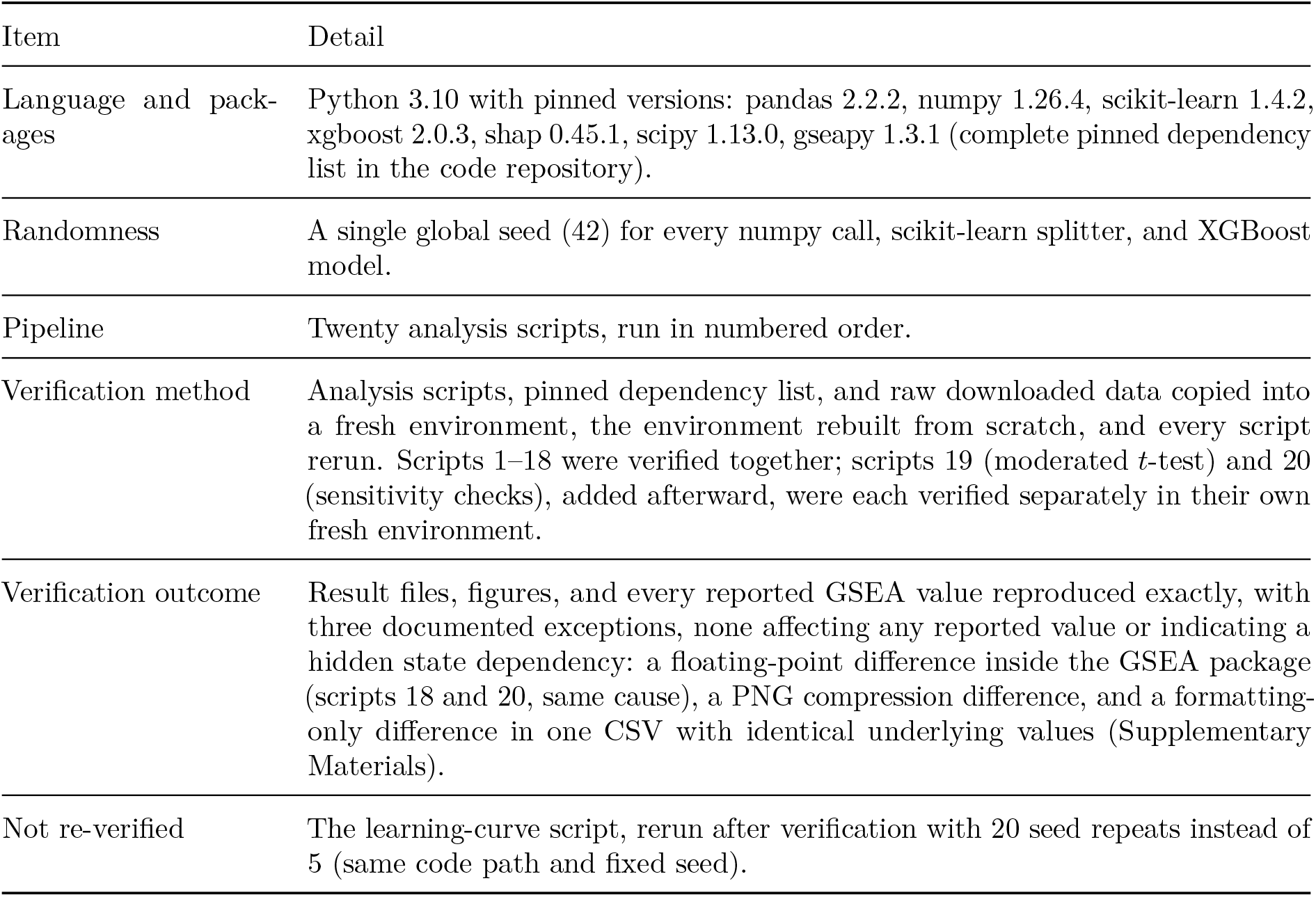
Software environment and reproducibility verification.

| Item | Detail |
| --- | --- |
| Language and packages | Python 3.10 with pinned versions: pandas 2.2.2, numpy 1.26.4, scikit-learn 1.4.2, xgboost 2.0.3, shap 0.45.1, scipy 1.13.0, gseapy 1.3.1 (complete pinned dependency list in the code repository). |
| Randomness | A single global seed (42) for every numpy call, scikit-learn splitter, and XGBoost model. |
| Pipeline | Twenty analysis scripts, run in numbered order. |
| Verification method | Analysis scripts, pinned dependency list, and raw downloaded data copied into a fresh environment, the environment rebuilt from scratch, and every script rerun. Scripts 1–18 were verified together; scripts 19 (moderated *t*-test) and 20 (sensitivity checks), added afterward, were each verified separately in their own fresh environment. |
| Verification outcome | Result files, figures, and every reported GSEA value reproduced exactly, with three documented exceptions, none affecting any reported value or indicating a hidden state dependency: a floating-point difference inside the GSEA package (scripts 18 and 20, same cause), a PNG compression difference, and a formatting-only difference in one CSV with identical underlying values (Supplementary Materials). |
| Not re-verified | The learning-curve script, rerun after verification with 20 seed repeats instead of 5 (same code path and fixed seed). |

Code is available at https://github.com/Salsabil-Ouni/Fibromyalgia-RNA-sequencing-Mac hine-Learning-Classifier-GSE221921. and, as a versioned, permanently archived snapshot of the exact commit used for this manuscript, at Zenodo (DOI: https://doi.org/10.5281/zenodo.23102139).

## 3 Results

### 3.1 Nested cross-validated performance

Both model families achieved similar performance: logistic regression ROC-AUC = 0.821 (95% CI 0.756–0.880), XGBoost ROC-AUC = 0.825 (95% CI 0.766–0.880) (Table 3). The paired bootstrap difference (XGBoost minus logistic regression) was +0.005 (95% CI −0.031 to 0.043), so there is no evidence of an advantage from the nonlinear model. Across five CV seeds, logistic regression AUC was 0.830 ± 0.011 (range 0.814–0.844), so the headline value is representative of split-to-split variability; the bootstrap intervals reflect sampling uncertainty of the pooled predictions and do not include split or model-selection variability.

**Table 3:** Nested cross-validation performance (pooled out-of-fold predictions, bootstrap 95% CI).

| Model | ROC-AUC | PR-AUC | Sensitivity | Specificity | Accuracy |
| --- | --- | --- | --- | --- | --- |
| Logistic regression (L1/L2) | 0.821 (0.756–0.880) | 0.841 (0.764–0.908) | 0.729 | 0.763 | 0.746 |
| XGBoost | 0.825 (0.766–0.880) | 0.864 (0.810–0.913) | 0.709 | 0.764 | 0.736 |

### 3.2 Permutation-label null baseline

Under permuted labels, using the same reduced hyperparameter grid as this null (Section 2.4), the pipeline reached a mean AUC of 0.501 ± 0.059 (range 0.363–0.623) across 100 reruns. The real model’s AUC under this identical reduced grid (0.827) falls far outside this distribution (empirical *p* = 0.0099; Figure 4), indicating the observed performance is not attributable to the feature-selection procedure alone operating on *n* ≪ *p* data with no true signal.

**Figure 4:**
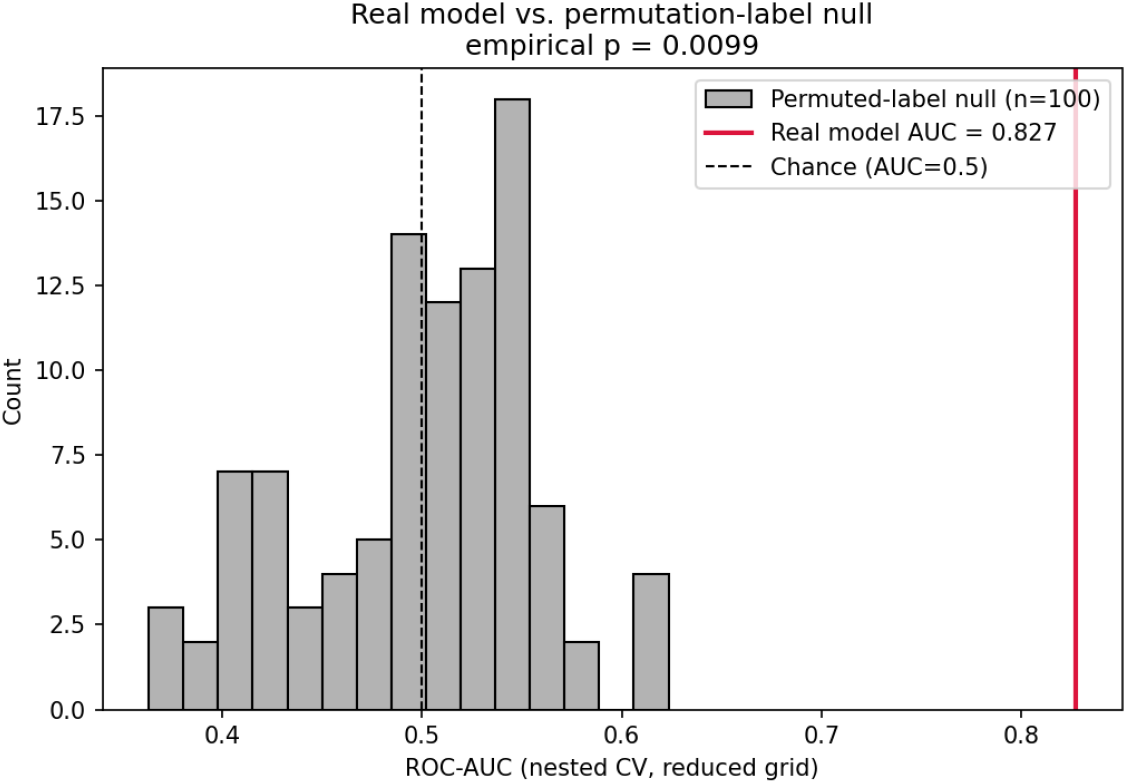
Real model AUC versus the permutation-label null distribution (100 reruns).

### 3.3 A sex confound in the cohort

Given the known sex imbalance of clinically-referred FM cohorts, the contribution of sex to classification was examined in detail. Sex was markedly imbalanced between groups: 91 of 96 FM patients (95%) were female, compared with 41 of 93 controls (44%) (*χ*^2^ = 55.3, *p* = 1.0 × 10^−13^). This distribution is consistent with the source publication and with the documented referral bias of clinic-based FM cohorts, which are approximately 95% female compared with 58–59% in population-based samples [9]; it is a known property of how FM cohorts are recruited, not a feature specific to this dataset.

The imbalance matters because a logistic regression classifier using sex alone, with no expression data, reaches ROC-AUC = 0.727 by 5-fold cross-validation (the group proportions imply about 0.75), close to the gene-expression model’s 0.821–0.825. How much expression signal survives once sex is accounted for was therefore checked using two structurally different approaches. Each has its own dedicated 100-rerun permutation-label null (Figure 5) rather than one borrowed from the whole-cohort analysis, since the smaller female-only cohort could plausibly have a different null distribution.

1. **Female-only subset.** Restricting the identical nested-CV pipeline to females (91 FM, 41 control; *n* = 132) yields ROC-AUC = 0.783 (95% CI 0.705–0.858), well above chance; its permutation null (*p* = 0.0099) rules out a small-*n* leakage artifact.
2. **Sex-chromosome gene exclusion.** Removing all 851 X- and Y-linked genes from the candidate feature pool while keeping the full, sex-imbalanced cohort yields ROC-AUC = 0.834 (95% CI 0.774–0.891), no decrease from the full-genome estimate (0.821; null *p* = 0.0099).

**Figure 5:**
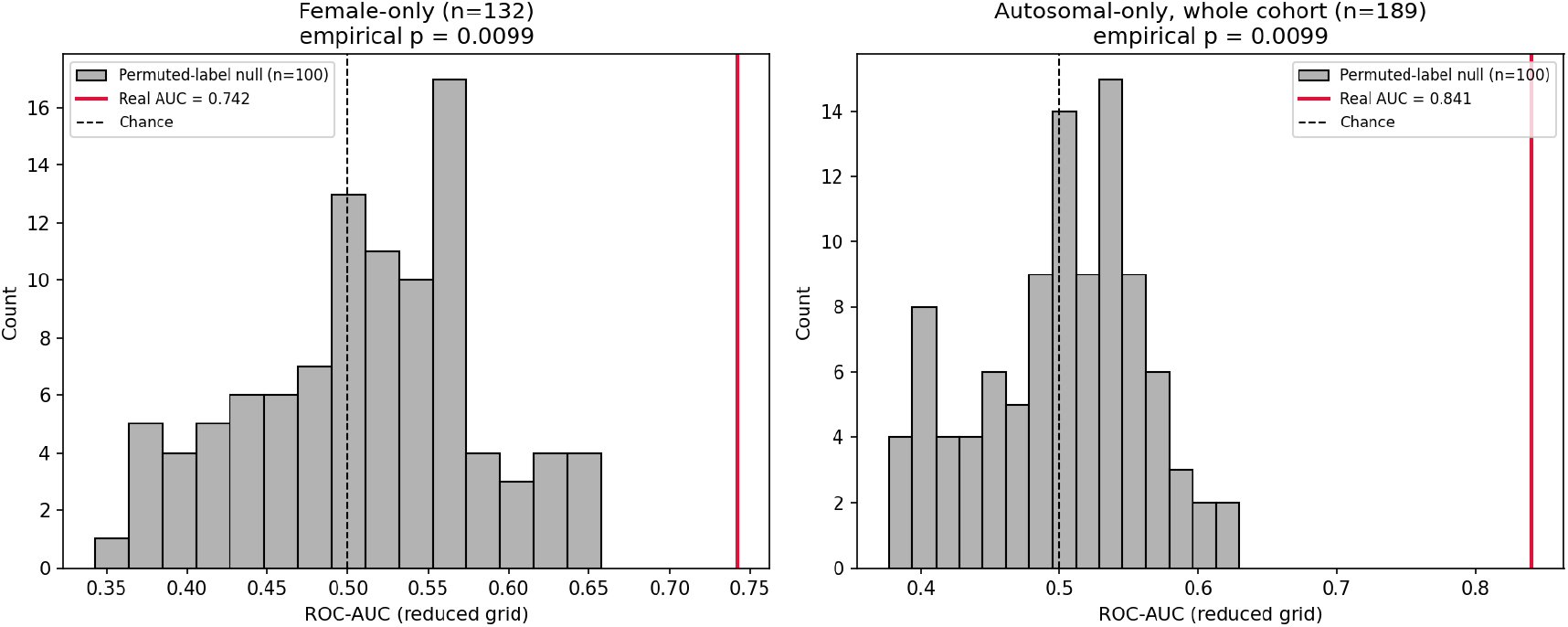
Permutation tests for the female-only model (left) and the model excluding X/Y-linked genes (right), 100 reruns each. Null means were 0.511 ± 0.071 and 0.500 ± 0.059; real-data AUCs under the same reduced grid were 0.742 and 0.841 (*p* = 0.0099 for both). The corresponding full-grid estimates are 0.783 and 0.834.

Because the female-only and whole-cohort AUCs have heavily overlapping confidence intervals, they were compared directly for the same 132 females using a paired bootstrap over subjects. The whole-cohort model scores slightly *lower* on these females (0.776) than the female-only model (0.783); the paired difference is −0.007 (95% CI −0.076 to +0.060, including zero), so training on the mixed-sex cohort does not measurably improve predictions for female patients.

The higher *pooled* AUC (0.821 versus 0.783 within females) is not driven by males (accuracy on the 57 males is only 0.592, with 5 male FM cases) but by a property of ROC-AUC: it counts every (FM, Control) pair, including cross-sex pairs that a sex-linked shift in predicted probability can rank correctly without within-sex discrimination. Across the 8,928 pairs, within-sex pairs are ranked correctly 76.4% of the time versus 86.7% for cross-sex pairs, reconstructing the pooled AUC exactly (0.8208); among females alone the rate is 77.6%, matching the whole-cohort model’s AUC on females (0.776), while the small male-male pairs (59.2%) lower the within-sex figure (Supplementary Table S4).

Removing the 851 X/Y-linked genes did not shrink the pooled AUC (0.834), which is consistent with this mechanism: the model’s sex correlation can arise from any gene whose expression differs by sex, including autosomal genes under hormonal or other indirect regulation, so it survives the exclusion. The two checks answer different questions: real disease signal survives once sex is removed structurally, and the classifier does not depend on direct chromosome dosage. The pooled metric remains partly shaped by autosomal sex-correlated expression, so the female-only estimate (0.783) is regarded here as the more defensible effect size.

Together, these checks show that the signal is not explained by sex, whether through X/Y-linked genes or the sex imbalance itself; Section 3.7 asks whether it reflects PBMC composition. Sex is not entirely absent from the gene ranking, however: 5 of the top 30 SHAP-ranked genes are X-linked (*ANOS1*, *KIF4A*, *TTC3P1*, *SAGE4P*, *FRMPD4* ), about four times the 3.9% share of X-linked genes in this dataset (binomial *p* = 0.005), though they do not dominate the list.

### 3.4 Calibration

Logistic regression’s predicted probabilities were reasonably well-calibrated (Brier score 0.171 vs. 0.250 for an uninformative base-rate-only baseline; Figure 6). XGBoost, despite matching AUC, was overconfident at high predicted probabilities (its ∼0.87 predicted-probability bin corresponded to an observed FM rate of only ∼0.58), indicating logistic regression’s probability outputs would be the safer choice for any downstream use that depends on calibrated probabilities rather than on discrimination or ranking alone. The two models remain statistically indistinguishable in discrimination (Section 3.1); this calibration gap does not bear on the explainability and pathway analyses in Sections 3.6–3.7, which depend on relative gene ranking rather than on predicted-probability values, but should be weighed by anyone considering either model for probability-based use beyond this paper.

**Figure 6:**
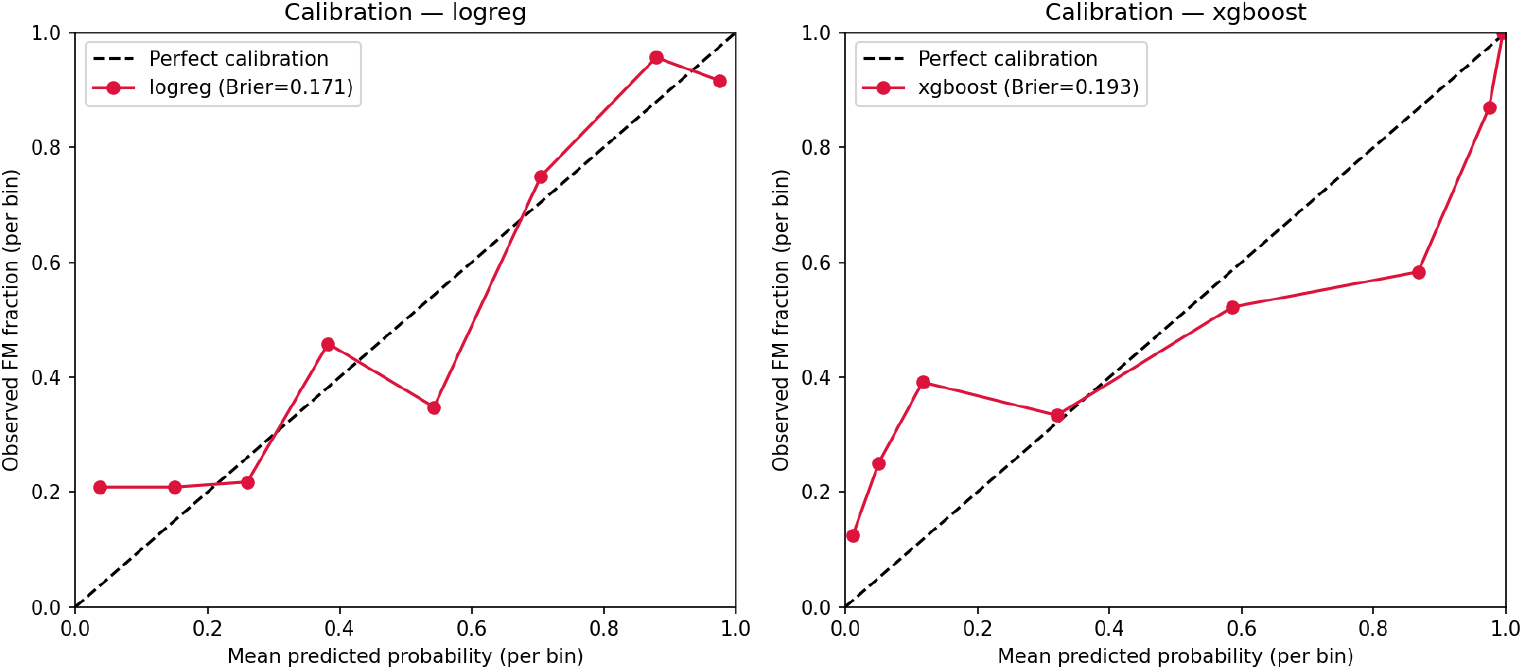
Calibration curves for logistic regression (left) and XGBoost (right).

### 3.5 Feature-selection stability, methodological robustness, and learning curve

Gene-level selection was noisy in aggregate across repeated folds (mean pairwise Jaccard overlap = 0.295 across 839 genes ever selected in 50 stability folds), as expected at *n* = 189. The genes the SHAP analysis ranked most important were stable, however: all top-30 SHAP genes were selected in ≥50% of folds (mean selection frequency 0.851 versus 0.172 across all genes; Supplementary Figure S1), so the reported genes are not fold-to-fold noise. Three further checks support the headline result (Supplementary Table S1). A Welch’s *t*-test selector shifts AUC by a real but modest +0.036 (paired 95% CI 0.006 to 0.069); the moderated *t*-test gives AUC = 0.822 (95% CI 0.757–0.883), a paired difference of +0.001 (−0.040 to 0.044) from the Mann-Whitney result; and four additional random seeds give AUC 0.838, 0.835, 0.844, 0.814 (std 0.011), showing that the headline is not an artifact of one split. Restricting the candidate gene pool to protein-coding genes only (16,834 / 21,915 genes; Section 2.5) gave AUC 0.845 (95% CI 0.786–0.900), a paired bootstrap difference of +0.024 from the full-genome result (95% CI −0.012 to 0.065, aligned sample-for-sample), so performance does not depend on the pseudogenes and noncoding transcripts present in the candidate pool, and if anything is nominally higher without them, though the difference is not distinguishable from zero. Performance (mean per-fold AUC, hence slightly higher than the pooled AUC) also rose gently with training-set size (20 seed repeats per point), from 0.806 at ∼46 training samples per fold to 0.842 at ∼152, without a clear plateau (Supplementary Figure S2); wide per-point standard deviations (0.06–0.08) make this trend suggestive rather than conclusive.

### 3.6 Top predictive genes and comparison to the original paper’s pathways

The XGBoost model, refit on the full dataset for interpretation only after the performance estimate in Section 3.1 was locked in, selected 50 features. The top-ranked genes by mean absolute SHAP value were *MIR1281*, *RPA2*, *FAM160A1* (current HGNC symbol *FHIP1A*), *TTC3P1*, *TMEM170B*, and *KIF4A* (Figures 7 and 8; the full 50-gene ranking, with current HGNC symbols noted for any gene renamed since the source dataset’s release, is given in Supplementary Table S2). Gene symbols throughout this manuscript are otherwise given as released in the source dataset, to match figures and supplementary tables generated directly from it.

**Figure 7:**
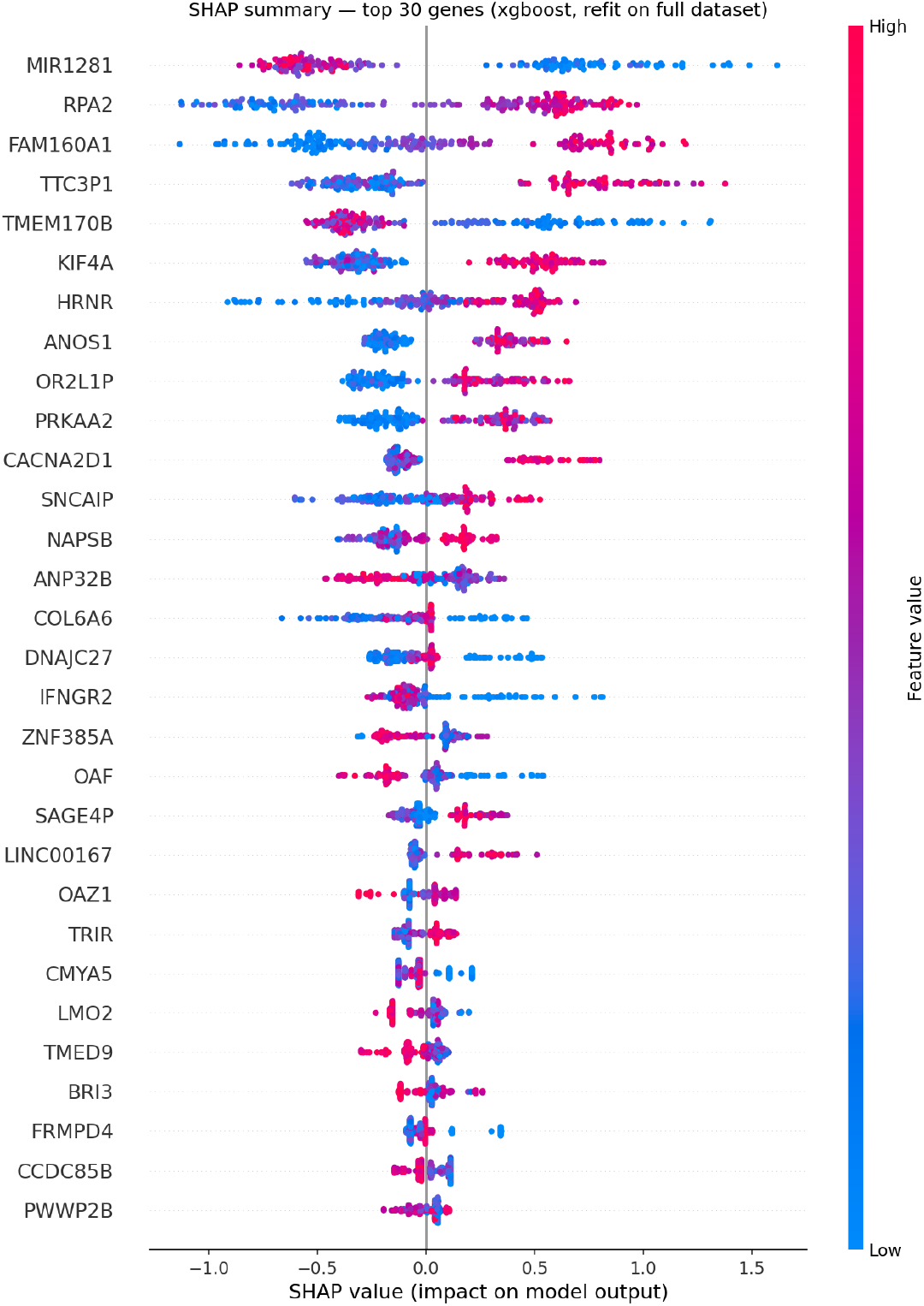
SHAP summary plot for the top 30 genes (XGBoost).

**Figure 8:**
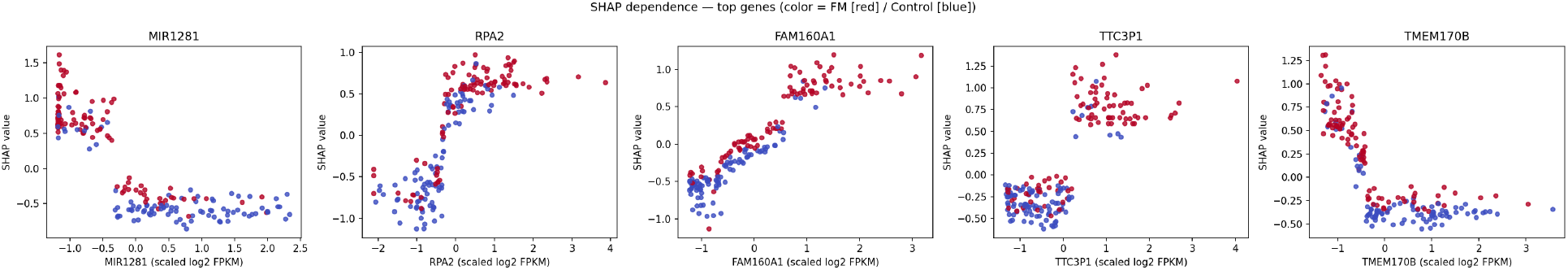
SHAP dependence plots for the top 5 genes, colored by FM/Control status.

XGBoost was used for this analysis because it had the nominally higher nested-CV AUC (Section 3.1), though that difference is not statistically significant, and logistic regression is independently known to be better-calibrated (Section 3.4). To make this model choice auditable rather than assumed, the identical SHAP analysis was repeated for logistic regression, refit and explained independently (LinearExplainer, full 50-gene ranking in Supplementary Table S2b). The two models’ top genes agree closely despite using unrelated mechanisms (coefficient-weighted linear SHAP vs. tree-ensemble SHAP): 8 of the top 10, 19 of the top 30, and all 50 of the top 50 genes are shared between the two rankings. This agreement, not the small and non-significant AUC difference, is the stronger justification for treating the reported gene ranking as a property of the expression data rather than of the particular model family used to surface it.

Preranked GSEA across the ranked list of 21,374 genes found the FM2 (CLEAR/lysosomal) and FM3 (interferon/JAK-STAT/death-receptor) gene sets significantly enriched, and FM1 (ECM/RhoGDI/collagen/ GP6) not (Table 5, Figure 9). Both enriched sets have negative normalized enrichment scores, meaning their genes sit toward the lower-expressed end of the FM-versus-control ranking, the reverse of the increased CLEAR/lysosomal expression reported for the original FM2 subgroup (discussed in Section 4). The enrichment persists in females alone (FM2 NES −2.56, FM3 NES −2.10, both FDR *q <* 0.001), so it is not a sex artifact, but it disappears after adjustment for cell-composition proxies (Section 3.7).

**Figure 9:**
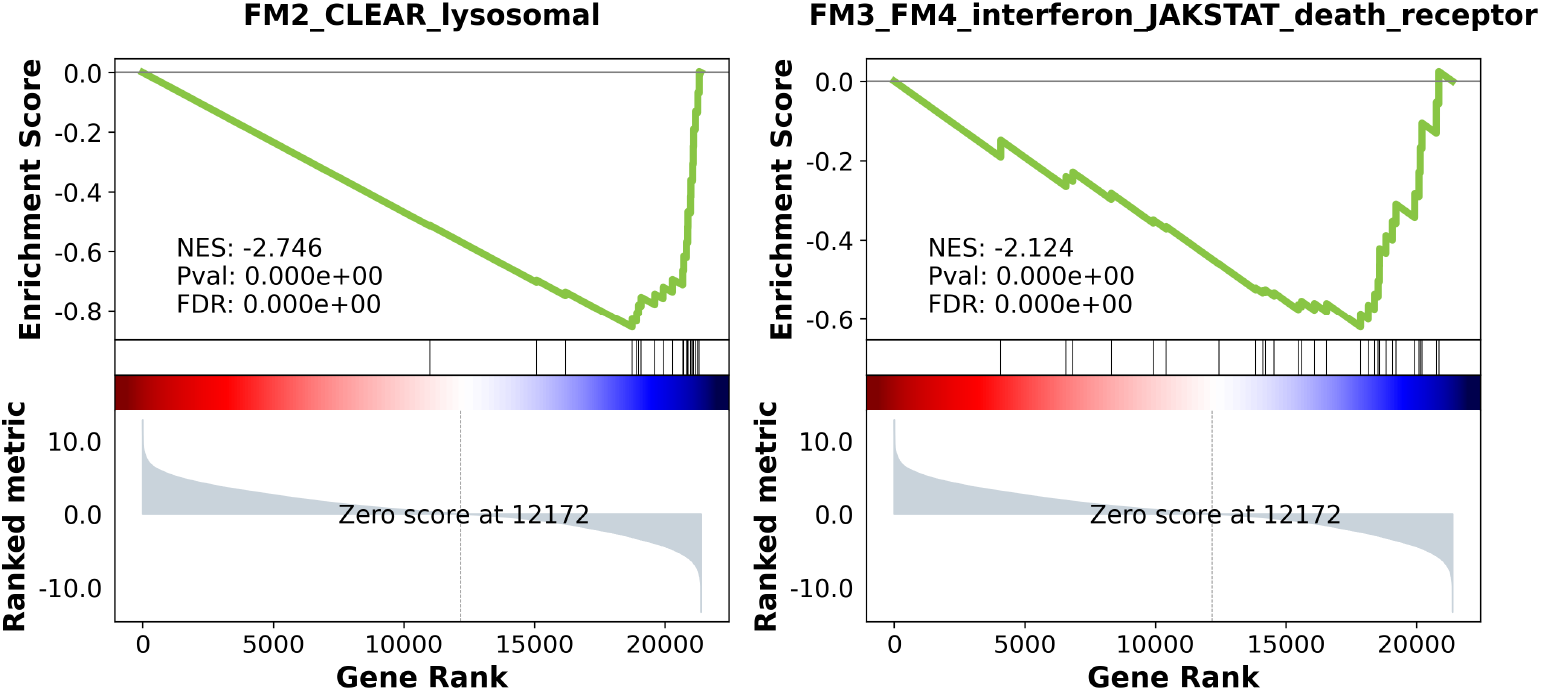
GSEA enrichment plots for the FM2 (CLEAR/lysosomal, left) and FM3 (interferon/JAK-STAT/death-receptor, right) gene sets. The corresponding plot for the non-significant FM1 gene set is given in Supplementary Figure S3.

**Table 4:** Confound checks: cohort composition and classifier performance under each condition.

| Condition | $n$ (FM / Control) | % female (FM / Control) | ROC-AUC (95% CI) |
| --- | --- | --- | --- |
| Sex only, no gene expression | 189 (96 / 93) | 95% / 44% | 0.727 (0.648–0.798)* |
| Whole cohort, all genes (Table 3) | 189 (96 / 93) | 95% / 44% | 0.821 (0.756–0.880) |
| Whole cohort, X/Y genes excluded | 189 (96 / 93) | 95% / 44% | 0.834 (0.774–0.891) |
| Female-only subset | 132 (91 / 41) | 100% / 100% | 0.783 (0.705–0.858) |
| Whole-cohort model, evaluated on females only | 132 (91 / 41)<sup>†</sup> | 100% / 100% | 0.776 (†) |
| Composition proxy only, whole cohort | 189 (96 / 93) | 95% / 44% | 0.755 (0.683–0.823)<sup>‡</sup> |
| Composition proxy only, females | 132 (91 / 41) | 100% / 100% | 0.712 (0.609–0.801)<sup>‡</sup> |

\*5-fold cross-validation only (single-feature logistic regression, outside the nested-CV framework), so not directly comparable in rigor to the nested-CV rows. <sup>†</sup>Not an independent model: predictions of the whole-cohort model restricted to the 132 females, used for the paired comparison reported in the text (no separate CI).
<sup>‡</sup>Logistic regression on four marker-based composition scores, 5-fold CV (Section 3.7). All nested-CV rows (full genome, X/Y-excluded, female-only) use the full hyperparameter grid from Section 2.3, including both L1 and L2 penalties; the permutation-null distributions each row is tested against (Section 2.4, Figure 5) use a reduced grid, L2 penalty only, for computational tractability across 100 reruns, so the real-AUC values entering each null comparison differ slightly from the headline nested-CV AUCs reported here.

**Table 5:** Preranked GSEA results for the three original subtype gene sets.

| Pathway | Leading edge / set size | ES | NES | $p$-value | FDR $q$ |
| --- | --- | --- | --- | --- | --- |
| FM1 (ECM/RhoGDI/collagen/GP6) | 14 / 35 | $-0.368$ | $-1.281$ | 0.104 | 0.120 |
| FM2 (CLEAR/lysosomal) | 20 / 23 | $-0.856$ | $-2.746$ | $< 0.001$ | $< 0.001$ |
| FM3 (interferon/JAK-STAT/death-receptor) | 15 / 30 | $-0.621$ | $-2.124$ | $< 0.001$ | $< 0.001$ |

ES: enrichment score. NES: normalized enrichment score. Leading edge: genes contributing to the peak enrichment score, out of the genes in the set. $p$ -values from 1,000 gene-set permutations (0 of 1,000 permuted sets reached as extreme an enrichment score for FM2 and FM3, hence $p < 0.001$ rather than 0). FDR $q$ : Benjamini-Hochberg-adjusted across the three pathways tested. The same analysis repeated in females only, and after adjusting for cell-composition proxies, is given in Supplementary Table S7.

A secondary hypergeometric test of overlap between the top-*N* SHAP genes and the same gene sets was also run; it is uninformative by construction at this set size (chance predicts only 0.02–0.08 overlapping genes, so zero overlap is the null expectation regardless of any true enrichment) and is reported only in Supplementary Table S3, not interpreted further here.

### 3.7 PBMC cell composition as a confound

Marker-based composition scores (monocyte, T, B, and NK cell; 20 genes in total) differed between groups in females for monocytes, which were lower in FM (mean score −0.40 versus +0.22; Mann-Whitney *p* = 0.0002, surviving Bonferroni correction across the four scores tested, adjusted threshold 0.0125). The remaining three scores showed smaller, non-significant group differences in the same direction as monocytes (T cell: −0.15 vs. +0.14, *p* = 0.10; B cell: −0.08 vs. +0.02, *p* = 0.40; NK cell: −0.01 vs. −0.08, *p* = 0.64); with only 91 FM and 41 control females, these comparisons are underpowered to detect effects smaller than the monocyte one, so the non-significant results should be read as inconclusive rather than as evidence that T, B, and NK composition genuinely does not differ between groups. A classifier using the four scores alone reached AUC 0.755 (95% CI 0.683–0.823) in the whole cohort and 0.712 (0.609–0.801) in females (Table 4). The female-only expression model (0.783) exceeded the female composition-only baseline by 0.072 (paired 95% CI −0.025 to 0.175), a difference not distinguishable from zero. Adjusting expression for the four scores before the enrichment analysis removed the FM2 and FM3 enrichment (Table 5), consistent with a lower monocyte contribution in FM PBMC rather than pathway-specific dysregulation. These proxies are coarse, and residualizing also removes any disease signal correlated with composition, so this is a conservative test. It shows that a composition contribution to both the classifier and the enrichment result cannot be excluded, not that the signal is purely compositional.

## 4 Discussion

This study addressed two questions. The first was whether a rigorously validated supervised classifier can discriminate FM from controls on the basis of PBMC gene expression. Expression achieved a nested cross-validated AUC of approximately 0.82 (0.78 in females), exceeding the permutation null and remaining robust to the sex analyses (Sections 3.1–3.3). However, a crude cell-composition proxy reached 0.71 in females and the expression model did not significantly exceed it, so the signal may partly reflect PBMC composition, chiefly a lower monocyte signal in FM (Section 3.7). The second question was whether the pathways associated with FM status line up with those implicated by the original subtyping. Curated proxies for two of the three subtype signatures (FM2 and FM3) were enriched among genes expressed lower in FM (*p <* 0.001 each; FM1 *p* = 0.104), also in females alone, but not after composition adjustment; the classifier’s top genes do not overlap them, as chance predicts.

The reversal relative to the original FM2 subgroup, which showed increased CLEAR/lysosomal expression, has two non-exclusive explanations. A pooled comparison against control can return a net negative direction if only a minority of patients (the original FM2 subgroup, *n* = 30 of 96) show an increase. Independently, a lower monocyte contribution would lower these genes, since lysosomal/CLEAR-network and type I interferon-response transcripts are both enriched in monocytes relative to lymphocytes in reference immune-cell RNA-seq panels [16]. Separating these explanations requires the original subtype labels and either cell counts or a deconvolution reference, neither available here.

A lower monocyte contribution and altered interferon-pathway signal in PBMC are biologically consistent with the broader neuroimmune and glial-activation mechanisms proposed for nociplastic pain generally [1], in which circulating innate-immune cell populations and their cytokine output are hypothesized to interact with central pain-processing circuits rather than act as a purely peripheral marker of tissue damage. The same peripheral immune and interferon signatures have been proposed in endometriosis-associated nociplastic pain [4], the condition with which FM most frequently co-occurs and shares central-sensitization mechanisms [5]. This parallel is a plausible biological rationale for the monocyte and interferon-pathway findings here, but it cuts in both directions: it is equally consistent with a genuine FM-associated neuroimmune signal and with gynecological or hormonal confounding of exactly this immune axis (Limitations), and the present data cannot distinguish between these.

Testing against simple confound baselines is regarded here as the main methodological lesson of this reanalysis. Sex composition alone gives AUC 0.727, close to what the group proportions imply (about 0.75), and a 20-gene composition proxy gives 0.71–0.76. Classifiers of clinically referred FM cohorts should be benchmarked against such baselines.

The biological interpretation of the top-ranked genes remains limited. Several are pseudogenes or noncoding transcripts (*MIR1281*, *TTC3P1*, *OR2L1P*, *SAGE4P*, *NAPSB*), which are prone to multi-mapping artifacts in RNA-seq quantification; since raw reads are not available, such artifacts cannot be excluded for individual genes. Nevertheless, restricting the candidate pool to protein-coding genes did not reduce performance (Section 3.5), which argues against a mapping artifact as the main driver of discrimination. Medication represents an additional and probably underestimated source of variation. The source study excluded only patients who had received anti-inflammatory drugs, or who had another chronic disease, within three months of enrollment [6]; exclusion or washout criteria for the analgesics, gabapentinoids, and antidepressants commonly prescribed in FM were not reported. Notably, *CACNA2D1* encodes the *α*_2_*δ*-1 subunit targeted by pregabalin and gabapentin, making a drug effect on this gene plausible. As any top-ranked gene may reflect drug exposure rather than disease biology, the ranking is interpreted here as a description of the classifier rather than as evidence of mechanism.

### Limitations

- **Single cohort.** All estimates derive from one PBMC RNA-seq dataset, so independent replication is needed. An AUC of ∼0.8 supports research prioritization rather than clinical diagnosis.
- **Cell composition.** Composition was inferred from 20 marker genes, without cell counts or validation against an established method (e.g. CIBERSORTx or a reference panel such as [16]), so composition cannot be separated from expression changes, and the marker-based scores themselves are unvalidated proxies. The four composition comparisons in Section 3.7 were not corrected for multiple comparisons; only the monocyte difference would survive a Bonferroni correction, and that is the one the enrichment result depends on. The non-significant T-, B-, and NK-cell comparisons are additionally underpowered at this sample size and should not be read as evidence that composition is unaffected for these cell types, only that any such effect, if present, is smaller than the monocyte one.
- **Unmeasured confounders.** Age, sequencing batch, medication, and comorbidities are absent from the public metadata and could not be tested. Sex was testable and was addressed by two independent checks; a male-only analysis was not feasible, as only five male FM patients were included.
- **Gynecological and hormonal confounding.** Patient sex is severely imbalanced in this cohort (95% female among FM patients vs. 44% among controls, Section 3.3), and gynecological and chronic pelvic pain conditions, including endometriosis, adenomyosis, and chronic pelvic pain syndrome, are frequent comorbidities of FM that share central-sensitization mechanisms with it. Hormonal status, including hormonal therapies (progestins, GnRH analogues, combined oral contraceptives), menopausal status, and menstrual cycle phase, are all known to alter PBMC monocyte and interferon-pathway gene expression. None of this is recorded in the public metadata for this dataset, so the female-only AUC (0.783) and the monocyte and interferon-pathway findings (Sections 3.6–3.7) may partly reflect gynecological comorbidity or hormone exposure rather than FM status specifically, and this cannot be tested or excluded with the data available here.
- **Subtype and label information.** The original subtype and reclassification labels are not public (only 2 of the 5 reclassified samples could be identified by ID), so the analysis compares pooled FM against control across all 189 samples and cannot test the direction of effect within the FM2 subgroup or the effect of label noise.
- **Enrichment analysis.** At *n* = 189, about 27% of genes share tied rank statistics, and the gene sets are curated proxies rather than the original study’s lists.

## Conclusions and perspectives

In this cohort, PBMC gene expression discriminates FM patients from controls, but a substantial part of this signal is attributable to sex imbalance and to blood cell composition, chiefly a reduced monocyte contribution in FM. Three priorities emerge for future research:

1. Replication in an independent cohort with balanced sex distribution and recorded age, medication, hormonal status, gynecological comorbidity (including endometriosis), and measured blood cell counts.
2. Reanalysis using the original subtype labels, in order to distinguish a genuine pathway alteration from subtype dilution and compositional effects.
3. Joint investigation of FM and endometriosis within the same benchmarking framework, to determine whether the monocyte and interferon signals are shared, condition-specific, or hormonally driven.

More generally, blood-based classifiers for chronic pain conditions should systematically report their performance relative to sex-only and composition-only baselines.

## Supplementary Materials

Supplementary Figures S1–S3, Tables S1–S7, and methodological detail are provided in a companion document, distributed alongside this file.

## Data and Code Availability

All analysis scripts, results tables, and figures are available at https://github.com/Salsabi l-Ouni/Fibromyalgia-RNA-sequencing-Machine-Learning-Classifier-GSE221921., with a versioned, permanently archived snapshot of the exact commit used for this manuscript at Zenodo (DOI: https://doi.org/10.5281/zenodo.23102139). The analyzed dataset is public at NCBI GEO (accession GSE221921) and is not duplicated there; the repository documents the download details and checksums, and its numbered scripts, run in order, regenerate the processed data and every result. A full reproducibility log is included.

## Acknowledgments

The authors thank the patients and researchers behind GSE221921 for making their data publicly available. This work is dedicated to people living with fibromyalgia, a condition still too often met with disbelief and diagnostic delay. The authors hope that rigorous research with careful attention to confounders will help develop objective diagnostic tools and better clinical care.

## Ethics Statement

This study is a secondary reanalysis of de-identified, publicly available data (GEO accession GSE221921); no new human subjects data were collected. Participant recruitment, informed consent, and institutional ethics approval for the original data collection are described in the source publication [6].

## Abbreviations

AUC: area under the curve
CI: confidence interval
CLEAR: Coordinated Lysosomal Expression and Regulation
ECM: extracellular matrix
ES: enrichment score
FDR: false discovery rate
FM: fibromyalgia
FPKM: fragments per kilobase of transcript per million mapped reads
GEO: Gene Expression Omnibus
GSEA: gene set enrichment analysis
JAK-STAT: Janus kinase-signal transducer and activator of transcription
MES: Ministry of Higher Education and Scientific Research
NES: normalized enrichment score
NK: natural killer
PBMC: peripheral blood mononuclear cell
PCA: principal component analysis
PR-AUC: area under the precision-recall curve
RhoGDI: Rho GDP-dissociation inhibitor
RNA-seq: RNA sequencing
ROC-AUC: area under the receiver operating characteristic curve
SD: standard deviation
SHAP: SHapley Additive exPlanations
XGBoost: eXtreme Gradient Boosting.

