## Supplementary Materials for "Machine learning classification of fibromyalgia from blood transcriptomes: confounder benchmarking and implications for endometriosis-associated pain"

This document accompanies the main manuscript and provides full detail for items summarized in the main text: the feature-selection stability, methodological robustness, and learning-curve checks (Section 3.5); the complete SHAP-ranked gene list (Section 3.6); the enrichment plot for the non-significant FM1 gene set from the primary GSEA pathway analysis (Section 3.6); the complete secondary hypergeometric pathway cross-check results (Section 3.6); the pairwise decomposition of the pooled AUC (Section 3.3); the curated pathway gene sets (Section 2.6); the cell-composition and model-comparison sensitivity analyses (Sections 3.1 and 3.7); and the detailed explanation of the two minor, inconsequential exceptions noted in the clean-room reproducibility verification (Section 2.8). Throughout this document, a plain "Section X.Y" refers to the main manuscript; sections of this document are numbered S1–S11, as listed below.

#### Contents

S1. Feature-selection stability 2

S2. Methodological robustness checks 3

S3. Learning curve 4

S4. Full SHAP-ranked gene list 4

S5. Logistic-regression SHAP ranking and comparison to XGBoost 6

S6. GSEA enrichment plot for the non-significant FM1 gene set 6

S7. Full pathway cross-check results (secondary, hypergeometric) 7

S8. Pairwise decomposition of the pooled AUC 8

S9. Curated pathway gene sets 9

S10. Additional sensitivity checks 9

S11. Reproducibility verification detail 10

### S1. Feature-selection stability

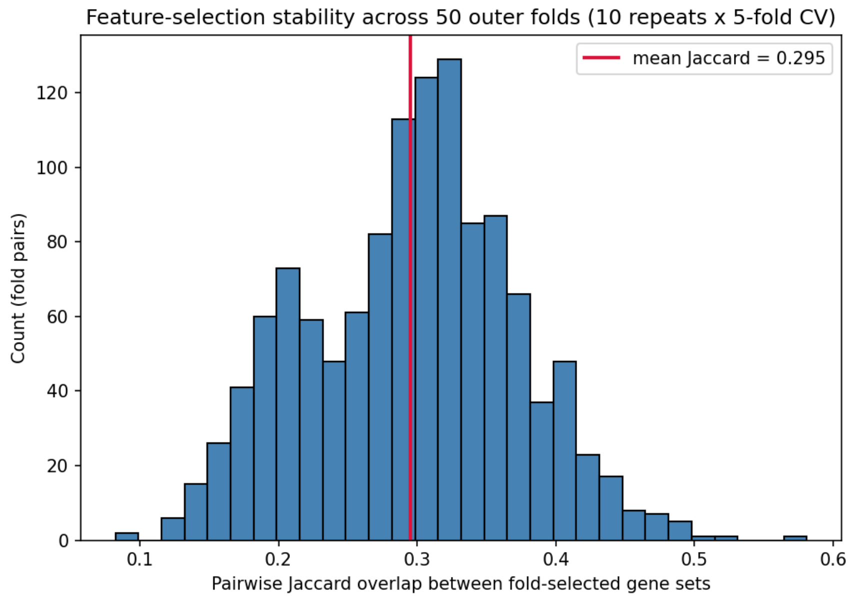


Figure S1: Distribution of pairwise Jaccard overlap between the gene sets selected in each of 50 outer-CV folds (10 repeats × 5-fold CV, logistic regression). Overall gene-level overlap is modest (mean = 0.295), but the top-30 SHAP-ranked genes are selected far more consistently than this aggregate figure suggests (main text, Section 3.5).

#### S2. Methodological robustness checks

Table S1: Feature-selection stability and methodological robustness checks (main text, Section 3.5).

| Check | Metric | Result |
|---|---|---|
| Feature-selection stability | Mean pairwise Jaccard overlap, all genes ever selected (839 genes, 50 folds) | 0.295 |
| Feature-selection stability | Mean selection frequency, top-30 SHAP genes | 0.851 |
| Feature-selection stability | Mean selection frequency, all genes ever selected | 0.172 |
| Feature-selection stability | Top-30 SHAP genes selected in $\geq$50% of folds | 30 / 30 |
| Univariate-test robustness | ROC-AUC, Mann-Whitney U selection (logistic regression) | 0.821 (0.756–0.880) |
| Univariate-test robustness | ROC-AUC, Welch's $t$-test selection (logistic regression) | 0.857 (0.801–0.907) |
| Univariate-test robustness | Paired bootstrap diff ($t$-test − Mann-Whitney) | +0.036 (0.006–0.069) |
| Univariate-test robustness | ROC-AUC, moderated $t$-test selection (logistic regression) | 0.822 (0.757–0.883) |
| Univariate-test robustness | Paired bootstrap diff (moderated $t$-test − Mann-Whitney) | +0.001 (−0.040–0.044) |
| Seed sensitivity (whole cohort, logistic regression, full grid) | AUC across 5 CV seeds (incl. headline seed 42) | mean 0.830, std 0.011, range [0.814, 0.844] |

Source: the corresponding result tables in the code repository accompanying the main manuscript. The moderated $t$-test is a from-scratch implementation of the Smyth (2004) empirical-Bayes algorithm underlying limma's eBayes function, validated on synthetic data before use (see the repository's moderated-test scoring module); it was not cross-checked against the limma R package itself.

#### S3. Learning curve

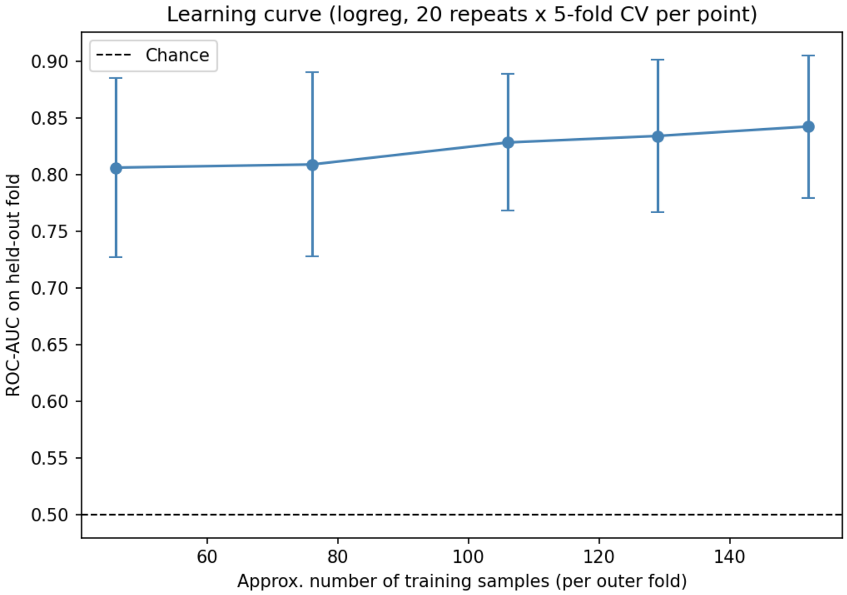


Figure S2: ROC-AUC on held-out folds as a function of approximate training-set size (main text, Section 3.5), holding test-fold composition fixed and subsampling only the training portion of each outer fold (logistic regression, 20 CV-seed repeats per point, error bars = standard deviation across 100 fold-evaluations per point).

#### S4. Full SHAP-ranked gene list

Table S2 lists all 50 genes selected by the final XGBoost model (refit on the full dataset for interpretation, main text Section 2.5 and 3.6), ranked by mean absolute SHAP value. The main text names only the top 6.

Table S2: Full SHAP feature importance ranking (main text, Section 3.6).

| Rank | Gene | Mean \|SHAP\| |
|---|---|---|
| 1 | MIR1281 | 0.6368 |
| 2 | RPA2 | 0.5900 |
| 3 | FAM160A1 | 0.4790 |
| 4 | TTC3P1 | 0.4781 |
| 5 | TMEM170B | 0.4389 |
| 6 | KIF4A | 0.4078 |
| 7 | HRNR | 0.3010 |
| 8 | ANOS1 | 0.2704 |
| 9 | OR2L1P | 0.2646 |
| 10 | PRKAA2 | 0.2630 |
| 11 | CACNA2D1 | 0.2184 |
| 12 | SNCAIP | 0.1929 |

| Rank | Gene | Mean \|SHAP\| |
|---|---|---|
| 13 | NAPSB | 0.1707 |
| 14 | ANP32B | 0.1704 |
| 15 | COL6A6 | 0.1660 |
| 16 | DNAJC27 | 0.1497 |
| 17 | IFNGR2 | 0.1422 |
| 18 | ZNF385A | 0.1362 |
| 19 | OAF | 0.1309 |
| 20 | SAGE4P | 0.1023 |
| 21 | LINC00167 | 0.1000 |
| 22 | OAZ1 | 0.0874 |
| 23 | TRIR | 0.0804 |
| 24 | CMYA5 | 0.0787 |
| 25 | LMO2 | 0.0737 |
| 26 | TMED9 | 0.0694 |
| 27 | BRI3 | 0.0646 |
| 28 | FRMPD4 | 0.0615 |
| 29 | CCDC85B | 0.0595 |
| 30 | PWWP2B | 0.0554 |
| 31 | TACC2 | 0.0457 |
| 32 | CLTB | 0.0412 |
| 33 | PPP5D1 | 0.0400 |
| 34 | TYROBP | 0.0393 |
| 35 | MRC1 | 0.0282 |
| 36 | IL1RAPL1 | 0.0271 |
| 37 | TRIM11 | 0.0244 |
| 38 | NINJ1 | 0.0210 |
| 39 | SLC26A5 | 0.0201 |
| 40 | MUC19 | 0.0135 |
| 41 | MOB2 | 0.0105 |
| 42 | DEDD2 | 0.0085 |
| 43 | MXRA5 | 0.0085 |
| 44 | LIPK | 0.0063 |
| 45 | C19orf38 | 0.0000 |
| 46 | TKT | 0.0000 |
| 47 | MRFAP1 | 0.0000 |
| 48 | CD68 | 0.0000 |
| 49 | TGFBI | 0.0000 |
| 50 | RNF130 | 0.0000 |

Source: the corresponding full SHAP-importance table in the code repository. The last 6 genes have mean |SHAP| = 0: they passed the univariate Mann-Whitney filter (so were among the 50 genes made available to the model) but were never used in any split of the refit XGBoost model, so they contribute nothing to its predictions despite being nominally “selected.” Gene symbols are given as released in the source dataset. Three have since been updated by HGNC: *FAM160A1* (rank 3) is now *FHIP1A*; *LINC00167* (rank 21) is now *PRDM10-DT*; *PPP5D1* (rank 33) is now *PPP5D1P*.

#### S5. Logistic-regression SHAP ranking and comparison to XGBoost

Table S3 gives the full logistic-regression SHAP ranking (main text, Section 3.6), computed independently from the XGBoost ranking in Table S2, as a check on whether the choice between the two nested-CV-tied model families changes which genes are identified as important. The refit model used an L1 penalty (selected by the same full hyperparameter grid as the nested-CV pipeline, Section 2.3), which drives most coefficients exactly to zero; only 12 of the 50 candidate genes retain nonzero weight and therefore nonzero SHAP value.

Table S3: Logistic-regression SHAP ranking (main text, Section 3.6).

| Rank | Gene | Mean \|SHAP\| |
|---|---|---|
| 1 | FAM160A1 (FHIP1A) | 0.3893 |
| 2 | MIR1281 | 0.2978 |
| 3 | RPA2 | 0.2904 |
| 4 | ANOS1 | 0.1941 |
| 5 | IFNGR2 | 0.1908 |
| 6 | TMEM170B | 0.1662 |
| 7 | KIF4A | 0.1486 |
| 8 | TTC3P1 | 0.1229 |
| 9 | OR2L1P | 0.0628 |
| 10 | PPP5D1 (PPP5D1P) | 0.0294 |
| 11 | CACNA2D1 | 0.0099 |
| 12 | MRC1 | 0.0032 |
| 13–50 | (remaining 38 genes) | 0.0000 |

Source: `results/shap_feature_importance_logreg.csv` in the code repository. Ranks 13–50 are the 38 genes the L1-penalized model assigned exactly zero coefficient; they are listed individually in the repository file. Top-$N$ overlap with the XGBoost ranking (Table S2): 8/10, 12/20, 19/30, 50/50, computed in `results/shap_model_overlap.csv`.

#### S6. GSEA enrichment plot for the non-significant FM1 gene set

Figure S3 shows the preranked GSEA enrichment plot for the FM1 (ECM/RhoGDI/ collagen/GP6) gene set (main text, Section 3.6, Table 4), which did not reach significance (NES $= -1.281$, $p = 0.104$, FDR $q = 0.120$). It is shown here for completeness alongside the two significantly enriched gene sets (FM2, FM3) plotted in the main text.

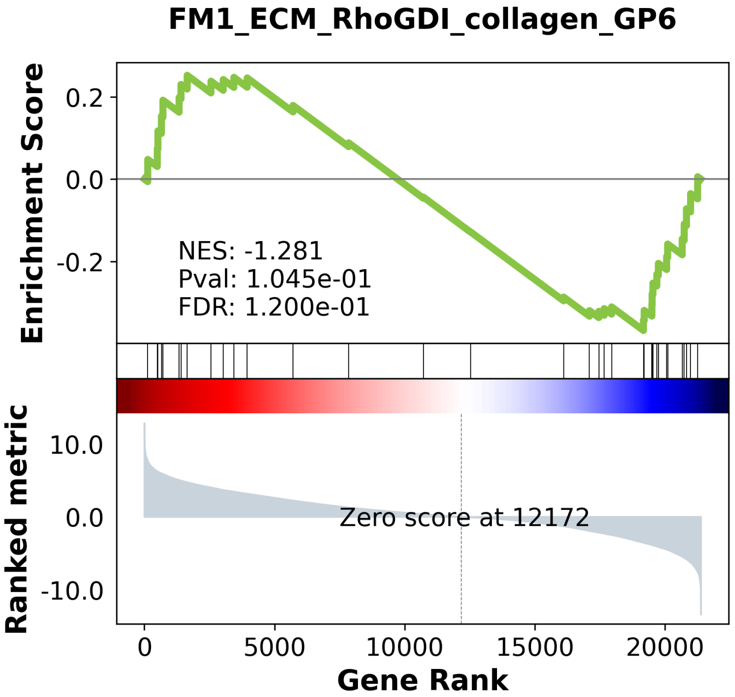


Figure S3: GSEA enrichment plot for the FM1 (ECM/RhoGDI/collagen/GP6) gene set (not significant).

#### S7. Full pathway cross-check results (secondary, hypergeometric)

Table S4 gives the complete hypergeometric overlap results across all four top-$N$ thresholds and all three pathway gene sets tested (main text, Section 3.6), including the minimum overlap count that would have reached $p < 0.05$ at each threshold. That minimum is not a measure of statistical power: by chance the expected overlap is only 0.02–0.08 genes (for example, $50 \times 35/21{,}915 = 0.08$ for FM1 at $N = 50$), so an overlap of zero is the null expectation and this check cannot distinguish overlap from its absence. It is a secondary check restricted to the classifier's own top-$N$ SHAP-ranked genes and answers a different question from the primary GSEA analysis (main text Table 4), which tests the same three gene sets for coordinated enrichment across the full transcriptome.

Table S4: Full secondary hypergeometric pathway cross-check results (main text, Section 3.6).

| Top-$N$ | Pathway | Overlap | $p$-value | Min. overlap for $p < 0.05$ |
|---|---|---|---|---|
| 20 | FM1 (ECM/RhoGDI/collagen/GP6) | 0 / 20 | 1.0 | 1 |
| 20 | FM2 (CLEAR/lysosomal) | 0 / 20 | 1.0 | 1 |
| 20 | FM3 (interferon/JAK-STAT/death-receptor) | 0 / 20 | 1.0 | 1 |
| 30 | FM1 (ECM/RhoGDI/collagen/GP6) | 0 / 30 | 1.0 | 1 |
| 30 | FM2 (CLEAR/lysosomal) | 0 / 30 | 1.0 | 1 |
| 30 | FM3 (interferon/JAK-STAT/death-receptor) | 0 / 30 | 1.0 | 1 |
| 50 | FM1 (ECM/RhoGDI/collagen/GP6) | 0 / 50 | 1.0 | 2 |
| 50 | FM2 (CLEAR/lysosomal) | 0 / 50 | 1.0 | 2 |
| 50 | FM3 (interferon/JAK-STAT/death-receptor) | 0 / 50 | 1.0 | 2 |
| 100* | FM1 (ECM/RhoGDI/collagen/GP6) | 0 / 50 | 1.0 | 2 |
| 100* | FM2 (CLEAR/lysosomal) | 0 / 50 | 1.0 | 2 |
| 100* | FM3 (interferon/JAK-STAT/death-receptor) | 0 / 50 | 1.0 | 2 |

*The $N = 100$ row is capped at the 50 genes actually selected by the final model (Section 3.6), so its overlap counts and results are identical to the $N = 50$ row. Pathway background gene counts (genes from each curated set present in the 21,915-gene expression universe): FM1 = 35, FM2 = 23, FM3 = 30. Source: the corresponding pathway cross-check table in the code repository.

#### S8. Pairwise decomposition of the pooled AUC

Table S5 decomposes the whole-cohort pooled ROC-AUC into its underlying (FM, Control) pairwise comparisons by pair type (main text, Section 3.3). ROC-AUC equals the fraction of (FM, Control) pairs in which the FM sample receives the higher predicted probability, with ties counted as one-half. The four pair types partition all $96 \times 93 = 8{,}928$ pairs, and their weighted average reproduces the pooled AUC exactly. Cross-sex pairs are ranked correctly more often than within-sex pairs, and the within-sex figure is pulled down by the small, noisy male-male component (only 5 male FM patients).

Table S5: Pairwise concordance of the whole-cohort model by pair type (main text, Section 3.3).

| Pair type | FM $n$ | Control $n$ | Pairs | Concordance |
|---|---|---|---|---|
| FM-female vs. Control-female | 91 | 41 | 3,731 | 0.7757 |
| FM-male vs. Control-male | 5 | 52 | 260 | 0.5923 |
| FM-female vs. Control-male (cross-sex) | 91 | 52 | 4,732 | 0.8842 |
| FM-male vs. Control-female (cross-sex) | 5 | 41 | 205 | 0.4683 |
| All within-sex pairs | | | 3,991 | 0.7637 |
| All cross-sex pairs | | | 4,937 | 0.8669 |
| All pairs (equals the pooled AUC) | 96 | 93 | 8,928 | 0.8208 |

#### S9. Curated pathway gene sets

Table S6 lists the members of the three curated gene sets, restricted to genes present in the expression matrix (main text, Section 2.6). The sets were curated from the literature; FM3 was corrected against the example genes named in the original paper. Curated genes absent from the expression matrix (GP6 for FM1; SQSTM1, GBA, and NEU1 for FM2; IFI27, IRF9, and RIPK3 for FM3) could not be tested.

Table S6: Members of the curated gene sets present in the expression matrix.

| Gene set | $n$ | Genes |
|---|---|---|
| FM1 (ECM/ RhoGDI/ collagen/ GP6) | 35 | *COL1A1, COL1A2, COL3A1, COL4A1, COL5A1, COL5A2, COL6A1, COL6A2, COL6A3, FN1, LAMA2, LAMB1, LAMC1, SPARC, THBS1, MMP2, MMP9, TIMP1, TIMP2, ITGA1, ITGA2, ITGB1, ARHGDIA, ARHGDIB, RHOA, RHOB, RHOC, RAC1, RAC2, CDC42, GP1BA, GP9, ITGA2B, ITGB3, VWF* |
| FM2 (CLEAR/ lysosomal) | 23 | *ASAH1, GAA, GNS, IFI30, PSAP, ATP6V0B, ATP6V0C, ATP6V0D1, ATP6V1B2, GABARAP, CTSA, CTSB, CTSD, CTSZ, HEXA, HEXB, NPC1, NPC2, TFEB, MCOLN1, LAMP1, LAMP2, GLA* |
| FM3 (interferon/ JAK-STAT/ death-receptor) | 30 | *IFI44, IFI44L, IFIT1, IFIT3, ISG15, ISG20, MX1, MX2, OAS1, OAS2, OAS3, STAT1, STAT2, JAK1, JAK2, TYK2, IRF7, IFNAR1, IFNAR2, USP18, CASP10, CASP8, FADD, TNFRSF10A, TNFRSF10B, RIPK1, MLKL, IKBKB, IKBKG, CHUK* |

One gene merits mention. *COL6A6*, a type VI collagen gene thematically related to the FM1 extracellular matrix signature, appears among the top 30 SHAP genes but was not in the curated FM1 set (which includes *COL6A1*, *COL6A2*, and *COL6A3*). The set was not expanded after seeing the classifier's output, to avoid hindsight bias.

#### S10. Additional sensitivity checks

Table S7 gives the detail behind the cell-composition and model-comparison analyses in main text Sections 3.1 and 3.7, and the full GSEA results for the same three gene sets repeated in females only and after composition adjustment (main text Section 3.6). Composition proxies are the mean standardized expression of marker genes: monocyte (*CD14, LYZ, CST3, FCN1, CSF1R, VCAN*), T cell (*CD3D, CD3E, CD2, IL7R, TRAC*), B cell (*MS4A1, CD79A, CD79B, CD19*), and NK cell (*NKG7, GNLY, KLRD1, NCAM1, KLRF1*); none belongs to a tested pathway gene set. They are coarse proxies, not a deconvolution.

Table S7: Composition proxies and paired model comparisons (main text, Sections 3.1 and 3.7).

| | | |
|---|---|---|
| *A. Composition scores in females (91 FM, 41 control): mean score, FM vs. control* | | |
| Monocyte | −0.40 vs. +0.22 | $p = 0.0002$ |
| T cell | −0.15 vs. +0.14 | $p = 0.10$ |
| B cell | −0.08 vs. +0.02 | $p = 0.40$ |
| NK cell | −0.01 vs. −0.08 | $p = 0.64$ |
| *B. Composition-only classifiers (logistic regression, 5-fold CV): ROC-AUC (95% CI)* | | |
| Monocyte score only, whole cohort | 0.731 (0.655–0.803) | |
| Four scores, whole cohort | 0.755 (0.683–0.823) | |
| Monocyte score only, females | 0.698 (0.595–0.796) | |
| Four scores, females | 0.712 (0.609–0.801) | |
| *C. Paired bootstrap AUC differences (95% CI)* | | |
| Female-only expression model (0.783) minus composition-only (0.712) | +0.072 (−0.025 to 0.175) | |
| XGBoost (0.825) minus logistic regression (0.821), whole cohort | +0.005 (−0.031 to 0.043) | |

Table S8: Preranked GSEA, females only and composition-adjusted (main text, Section 3.6).

| Pathway | NES (females only) | FDR $q$ | NES (composition-adj.) | FDR $q$ |
|---|---|---|---|---|
| FM1 (ECM/RhoGDI/collagen/GP6) | −1.166 | 0.230 | −0.714 | 0.905 |
| FM2 (CLEAR/lysosomal) | −2.563 | < 0.001 | +0.769 | 0.832 |
| FM3 (interferon/JAK-STAT/death-receptor) | −2.096 | < 0.001 | −1.072 | 0.653 |

Females only: 91 FM, 41 control, same method as main text Table 4. Composition-adj.: females only, expression residualized on the four composition scores in Table S7A before ranking and enrichment.

#### S11. Reproducibility verification detail

Main text Section 2.8 reports that a clean-room rerun of the complete pipeline reproduced every non-GSEA result file, every figure, and every reported GSEA value exactly, with three documented exceptions, none of which affects any value reported in the manuscript. All three are detailed here for completeness. (The learning-curve outputs were regenerated with more seed repeats after this verification and are not covered by it, as stated in the main text.)

First, one floating-point value in the GSEA results (the FM3 normalized enrichment score) differed from the original run in its 16th significant digit (−2.1237525659550203 versus −2.12375256595502). This is consistent with non-deterministic floating-point summation order in the GSEA package's internal parallelized permutation aggregation, a known property of that kind of parallel computation rather than a hidden state dependency in the analysis code itself; it has no effect on any value reported at the precision used anywhere in this work.

Second, the 9 Matplotlib-generated figures (all figures except the 4 GSEA enrichment plots) were pixel-identical between the original run and the clean-room rerun, verified by direct pixel-array comparison rather than by file checksum alone, but differed in their underlying PNG compression byte stream (both runs recorded the same embedded Matplotlib version and DPI metadata). This most likely reflects a difference in the zlib/libpng build linked by the image library between the two Python environments, not any difference in image content.

Third, when the twentieth script (sensitivity checks, added after the main verification) was independently re-verified, two of its two GSEA-based rows (the composition-adjusted FM2 and FM3 enrichment scores) differed from the original run in their 15th-16th significant digit, the same non-deterministic-summation cause as the first exception above. A third output file (composition proxy scores by sex) differed at the byte level but contained identical values on parsing, a formatting-only artifact of how the two environments serialized the same floats. The other four output files from this script, including the paired model-comparison results, were byte-for-byte identical.
